# Targeted blocking of gene splicing can dysregulate intron-embedded microRNAs

**DOI:** 10.1101/2024.12.04.626772

**Authors:** Md Hasan Ali, Wojciech Kwiatkowski, Piotr Kopeć, Naveen Nedunchezhian, Sebastian Pęcherz, Natalia Kowalewska, Barbara Gnutti, Dario Finazzi, Savani Anbalagan

## Abstract

ASOs (antisense oligonucleotides) are a promising therapeutic approach for suppression, induction of gene expression or the correction of aberrant splicing. Addressing whether ASOs targeting genes embedded with intronic noncoding RNAs (ncRNAs) affect the expression and function of intronic ncRNAs is of importance to the success of ASOs in clinical trials. While studying the development of the zebrafish posterior pituitary (neurohypophysis), an important neuroendocrine interface, we observed that an ASO targeting the splice site, in contrast to the one targeting the translation site of the gene *slit3*, disrupts neurohypophyseal axonal morphogenesis. In addition to altered *slit3* splicing, we also observed an increase in the expression of *slit3* and *slit3* intron-embedded primary *mir218a-*1 transcripts. The ASO-induced phenotype was not observed when mature *mir218a-1* was blocked by an ASO or in *mir218a-1*^-/-^ mutants. In addition, we also found that previously reported phenotypes due to ASOs targeting the splice site of *pank2* and *dnm2a* were partially rescued when the mature *mir103* and *mir199-5p* embedded in their introns, respectively, were blocked by ASOs. Our observation that ASOs targeting splice sites can affect intronic microRNA expression and function warrants further validation for other classes of ncRNAs. In addition, the idiosyncratic phenotypes when using translation and splice-blocking ASOs can be potentially used as a marker to identify the role of intronic ncRNAs.

## INTRODUCTION

Antisense oligonucleotides (ASOs) are one of the promising therapeutic options for monogenic disorders. Approximately 3000 human protein-coding genes are predicted to be haploinsufficient and a quarter of them are associated with diseases (Lek et al. 2016; Punta et al. 2015). In addition, splice-modulating ASOs are also under consideration for developing patient-specific ASOs (Kim et al. 2019; Aartsma-Rus et al. 2023). However, some ASOs can differentially affect mRNA vs pre-mRNA levels (Liang et al. 2017). Generally, when testing the potential of candidate ASOs *in vitro* and in animal models, the focus is largely on the rescue of the affected protein function and/or the disease phenotype. However, relatively few studies have addressed if the intronic non-coding RNAs (ncRNAs) in the targeted genes are affected by the host gene-targeting ASOs and the potential role of such ncRNAs in failure of ASOs in clinical trials (Kuhnert et al. 2008; Lacko et al. 2017; Fish et al. 2008).

In some tissues such as the brain or liver, intronic RNA can constitute a major proportion of the total cellular RNA (Nakaya et al. 2007; Kapranov et al. 2010; Wierzbicki et al. 2021; Mattick et al. 2023). Intronic ncRNAs types includes small nucleolar RNAs (snoRNAs), microRNAs (miRNAs), long non-coding RNAs (lncRNAs), stable intronic sequence RNAs (sisRNAs), and some genes can contain multiple types of ncRNAs (Wierzbicki et al. 2021; Mattick et al. 2023; Chan and Pek 2019; Statello et al. 2021). ncRNAs are implicated in various molecular biological processes, human diseases and are even considered as drug targets (Di Leva et al. 2014; Nemeth et al. 2024; Matsui and Corey 2017). Moreover, in some evolutionary conserved mammalian genes, there is strong divergence in the splicing pattern and length of the introns (Barbosa-Morais et al. 2012; Schaefke et al. 2018; McCoy and Fire 2020; William Roy and Gilbert 2006; Batzoglou et al. 2000; Roy et al. 2003; Pozzoli et al. 2007). Hence, any ASOs targeting such conserved yet “intronically” divergent genes in mouse may only offer a partial truth of how the host-gene-targeted ASOs can affect intronic ncRNAs in a human cell.

One of the well-studied ncRNA are the miRNAs and approximately 40-50% of all the miRNAs in human, mice and chicken genome are of intronic origin (Kim and Kim 2007; Carthew and Sontheimer 2009; Godnic et al. 2013). And this number is likely even higher as using DROSHA mutant proteins in *in vitro* studies, 75% of the human primary (pri-) miRNAs were shown to be intronic (Chang et al. 2015). In metazoans, majority of the host genes and their intronic pri-miRNAs are transcriptionally linked and coordinately expressed, while some intronic miRNA can be transcribed independently (Baskerville and Bartel 2005; Kim and Kim 2007; Rodriguez et al. 2004; Ramalingam et al. 2014; Biasiolo et al. 2011). Similar independent transcriptional units act as the major miRNA biogenesis mechanism in plants (Axtell et al. 2011; Gonzalo et al. 2022). The pri-miRNA transcripts are sequentially processed by the nuclear RNA-binding protein DGCR8/DROSHA and the cytoplasmic protein DICER1 to generate precursor and mature miRNAs, respectively (Bofill-De Ros and Vang Ørom 2024).

In addition, pre-miRNA can also be generated independent of DROSHA-mediated cleavage from small debranched introns (mirtrons) or short Polymerase II transcripts (Ruby et al. 2007; Xie et al. 2013). In human genome, there are 100s of such mirtrons (Da Fonseca et al. 2019; Wen et al. 2015). Some intronic miRNAs can target their host genes and function in a positive- or negative- feedback loop. (Hinske et al. 2010; Lutter et al. 2010) Several of monogenic human disease-linked genes are embedded with intronic miRNAs (Casar Tena et al. 2019; Zizioli et al. 2016). In some host gene-intronic miRNA pairs, the interpretation of the role of the host gene vs intronic miRNA required different tools, reagents and model organisms (Kuhnert et al. 2008; Lacko et al. 2017; Fish et al. 2008). Hence, if ASOs targeting gene splicing must succeed in clinical trials, it is critical to understand if blocking the splicing of host genes affects the intronic miRNA expression and function.

Apart from the potential of ASOs in therapeutics, ASOs were also widely used in developmental biology studies and are still in use. For instance, morpholino-based ASOs were extensively used in research involving zebrafish embryos with close to 3000 articles to study a wide variety of tissue and organ development and model human diseases. Several studies have shown that ASO-based phenotypes in zebrafish embryos can be either reproduced in the zebrafish or mice mutants. For instance, like mice LIM homeodomain transcription factor Islet1 knockout, ASO-based *islet1* knockdown in zebrafish leads to loss of spinal cord motoneurons (Hutchinson and Eisen 2006; Pfaff et al. 1996). Likewise, ASO-based *fgf8* knockdown has been demonstrated to phenocopy zebrafish *acerebellar* mutants *(fgf8* ^mut/mut^) and lack cerebellum and the midbrain-hindbrain boundary (Araki and Brand 2001).

However, despite the fact that several ASOs can phenocopy the mutants, some ASO’s can cause non-specific phenotypes that cannot be reproduced by mutants (Wright et al. 2004; Kok et al. 2015; Eisen and Smith 2008). We observed a similar discrepancy in phenotypes in our results of *pank2* splice-blocking ASO injected zebrafish embryos and *pank2*^-/-^ mutant embryos (Zizioli et al. 2016; Khatri et al. 2020; Mignani et al. 2022). Generally, the idiosyncrasy in the strong phenotypes arising from ASO-injected embryos versus lack of a phenotype in zebrafish mutants are largely attributed to Tp53 (P53)- based off target effects (15-20% of all ASO experiments) (Robu et al. 2007; Stainier et al. 2017; Lai et al. 2019; Kok et al. 2015; Eisen and Smith 2008). In our case, the injection of the *pank2*-specific ASO in *pank2*^-/-^ embryos, did not result in any phenotype, thus excluding this hypothesis (Khatri et al. 2020). Alternatively, the discrepancy can be due to genetic compensation or lack of maternal zygotic genetic background (Robu et al. 2007; Stainier et al. 2017). Even in the current guidelines for morpholinos use in zebrafish, the possible indirect effects of the intronic ncRNAs in the targeted gene have not been addressed (Stainier et al. 2017). However, the fact that off-target phenotypes arising from blocking the splicing of a single gene are dependent on the ASO target sequence suggests additional roles due to the aberrant products of unspliced transcripts (Robu et al. 2007).

Whether ASOs targeting the splicing of genes embedded with intronic miRNA inadvertently affect embedded pri-miRNA transcripts expression and its mature miRNA function is still under debate. We addressed this question in three zebrafish genes *slit3*, *pank2* and *dnm2a* that contain the intronic *mir218a-1*, *mir103* and *mir199-2a* respectively. In this study, using zebrafish embryos as vertebrate model, we show that the phenotypes due to splice-blocking ASO targeting at least 3 of the intronic miRNA-embedded host genes *slit3*, *pank2* and *dnm2a* can be partially or completely attributed to the increased intronic miRNA-encoding transcripts and their functional dysregulation.

## RESULTS AND DISCUSSION

### *slit3* splice-blocking ASO upregulates intronic miRNA transcript and perturbs pituitary axon development

Gene-embedded miRNAs are also in the introns of zebrafish protein-coding genes. Firstly, we performed a data-mining approach to identify all the zebrafish genes with intronic miRNAs as listed in the NCBI RefSeq annotated database. Out of 27,158 protein-coding genes, 66 genes contain intronic miRNAs and the majority of them contain 1 or 2 intronic miRNAs (**Fig. 1A-B** and **Table S1**). GO (gene ontology) term analysis revealed that intronic miRNA-containing genes were associated with functions such as developmental morphogenesis, calcium ion regulation, and other processes. (**Fig. 1C-D, Table S2**). During our datamining analysis, we also found 10 and 65 protein-coding genes with intronic snRNAs and snoRNAs, respectively (**Supplementary Fig. 1A-C, Table S3** and **S4)**. At least for these three classes of ncRNAs, we did not find any host genes that contain more than one class of ncRNA (data not shown).

**Figure 1.**
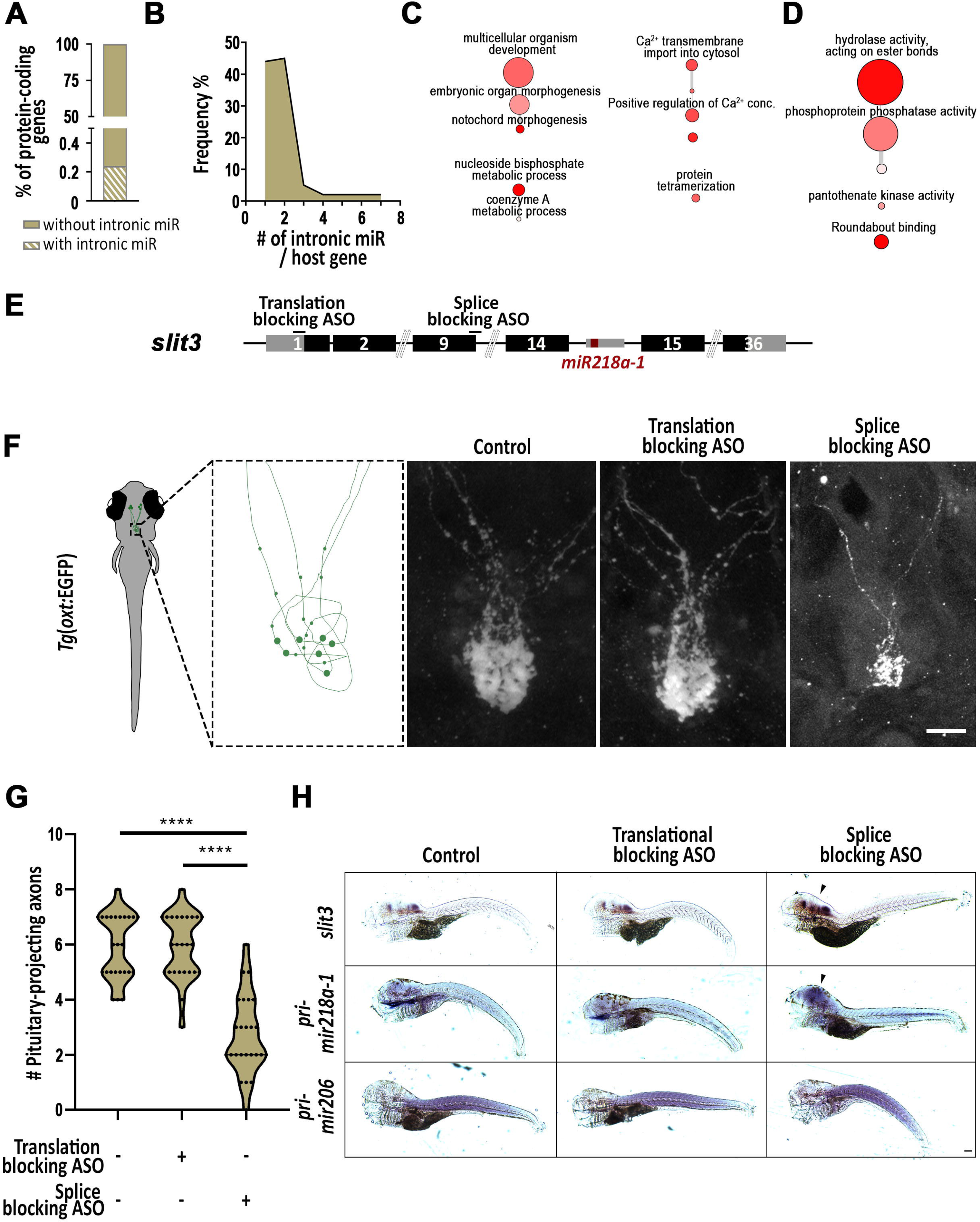
ASO targeting *slit3* splicing upregulate *pri-mir218a-1*. (**A**) Graph showing % of zebrafish protein coding genes with and without intronic miRNA. (**B**) Graph showing the frequency % and number of miRNA per miRNA-embedded host gene. (**C-D**) Figure showing the enriched biological process (**C**) and molecular function (**D**) GO terms in host genes. The color and size of the bubble correspond to the P value and LogSize value of the GO Term, respectively. (**E**) Scheme of *slit3 gene*. ASOs designed to target *slit3* translation and splicing are denoted by black lines and labelled. (**F**) Scheme describing the pituitary axons in (dpf) transgenic reporter Tg(*oxt*:EGFP) larvae and representative images showing whole-mount confocal microscope imaging of pituitary of 5 dpf transgenic Tg(*oxt*:EGFP) zebrafish following immunostaining with anti-EGFP. Images are from control embryos and following microinjection of *slit3* translation- or splice-blocking ASOs. Scale 20 µm. (**G**) Violin plot showing the number of pituitary-projecting OXT axons in 5 dpf control (n = 15) vs *slit3* translation-blocking ASO injected embryos (n = 14) and *slit3* splice-blocking ASO injected embryos (****p<0.001; Kruskal-Wallis test). Analysis were performed from 3 independent injections. (**H**) Representative images showing the expression of *slit3*, *pri-mir218a-1* and *pri-mir206* observed by digoxigenin-labeled probes in 5 dpf control embryos and following microinjection of translation- or splice-blocking ASO against *slit3*. Scale 100 µm.

Next, we wanted to narrow down few genes in which we could investigate if targeting the gene will affect the intronic miRNA embedded in the gene. Towards this, we compared the 66 miRNA-embedded genes with the zebrafish posterior pituitary glial cell-enriched transcriptome (Anbalagan et al. 2018). We identified 30 intronic miRNA-embedded genes with a minimum normalized expression count of 10 that were enriched in the glial transcriptome genes (**Table S5**). Out of these 30 genes, we focused on *slit3*, an evolutionarily conserved axonal guidance protein-coding gene (Blockus and Chedotal 2016)*. slit3* contains *mir218a-1* in the intron between exon 14 and 15 that has been previously reported to target Slit receptor-coding *robo* genes (**Fig. 1E**) (Fish et al. 2011).

To investigate if targeting *slit3* with a splice-blocking ASO will differently affect pituitary axons, we performed *slit3* knockdown experiments in zebrafish embryos with two different ASOs (morpholinos). The first ASO was a previously validated *slit3* translation-blocking ASO (Barresi et al. 2005). The second ASO was designed to block the *slit3* splicing by targeting the exon 9-intron 9 junction (**Fig. 1E**). We used the oxytocin neuron transgenic reporter line Tg(*oxt*:EGFP) to visualize and quantify the number of pituitary-projecting axons (**Fig. 1F**). In comparison to control and translation-blocking ASOs-injected larvae, we observed a reduced number of pituitary-projecting axons in the 5 days post fertilization (dpf) larvae that were injected with the splice-blocking ASO at one-cell stage (**Fig. 1F-G**). This phenotype was not due to oxytocin neuron deficits, as the number of oxytocin neurons in the neurosecretory preoptic area (NPO) was similar between control, translation- and splice-blocking ASO injected larvae (**Supplementary Fig. 1D-F, Video S1** and **S2**).

Since ASO-based phenotypes in zebrafish embryos can be also due to off-target effects mediated via Tp53, we tested if *tp53* knockdown can suppress the *slit3* splice-blocking ASO-induced phenotype (Robu et al. 2007). In comparison to control and *slit3* splice-blocking ASO injected embryos, co-injection of *tp53* translation-blocking ASO with *slit3* splice-blocking ASO did not result in any rescue in the number of pituitary-projecting axons (**Supplementary Fig. 1G**). These results suggest that *slit3* splicing-blocking ASO induced pituitary axonal defects are not due to TP53-based off-target effects that can be encountered in ASO studies.

We next tested if the two *slit3*-targeting ASO’s affect the expression of *slit3* and *slit3*-embedded primary *mir218a-1* transcripts. To observe primary *mir218a-1* transcript expression, we designed and synthesized a probe that span the entire primary *mir218a-1* and adjacent sequences (He et al. 2011). As a control for primary *mir218a-1* expression, we chose *pri-mir206* which has been previously reported to be expressed in the trunk muscles of zebrafish embryos (Kloosterman et al. 2006). In control samples, unlike *slit3* which has higher expression in forebrain and hindbrain, *pri-mir218a-1* expression was broader and we did not observe similar enrichment in the forebrain and hindbrain (**Fig. 1H**). In comparison to control larvae, we observed an increase in expression of *slit3* and *pri-mir218a-1*-coding transcripts in the splice-blocking ASO-injected samples. Unlike a strong increase in *pri-mir218a-1* transcripts, we did not observe a similar increase in the expression of *pri-mir206* in *slit3* splice-blocking ASO injected larvae (**Fig. 1H**). These results suggest that blocking of *slit3* splicing increases the expression of *slit3* and the *slit3* intron-embedded primary *mir218a-1* transcripts.

### *slit3* splice block ASO-induced phenotype is due to intronic *mir218a-1*

Splice-blocking ASOs can trigger exon skipping or intron retention in transcripts of the targeted gene (Eisen and Smith 2008). As expected, *slit3* splice-blocking ASO led to retention of the intron 9 (**Fig. 2A-B**). Since we observed the expression of misspliced *slit3* transcripts in splice-blocking ASO-injected embryos, we next tested whether the axonal phenotype was due to the potential truncated Slit3 protein or mature *mir218a-1* from misspliced *slit3* transcripts and the upregulated pri-*mir218a-1* transcripts, respectively. Toward this, we first performed ASO co-injection experiments. Specifically, we co-injected the *slit3* splice-blocking ASO with either *slit3* translation site-targeted or LNA-based *mir218a*-targeted ASO (**Fig. 2C**). Only in the samples in which *slit3* splice-blocking ASO was co-injected with the *mir218a*-targeting ASO, we observed a rescue in the number of pituitary-projecting axons (**Fig. 2D** and **E**).

**Figure 2.**
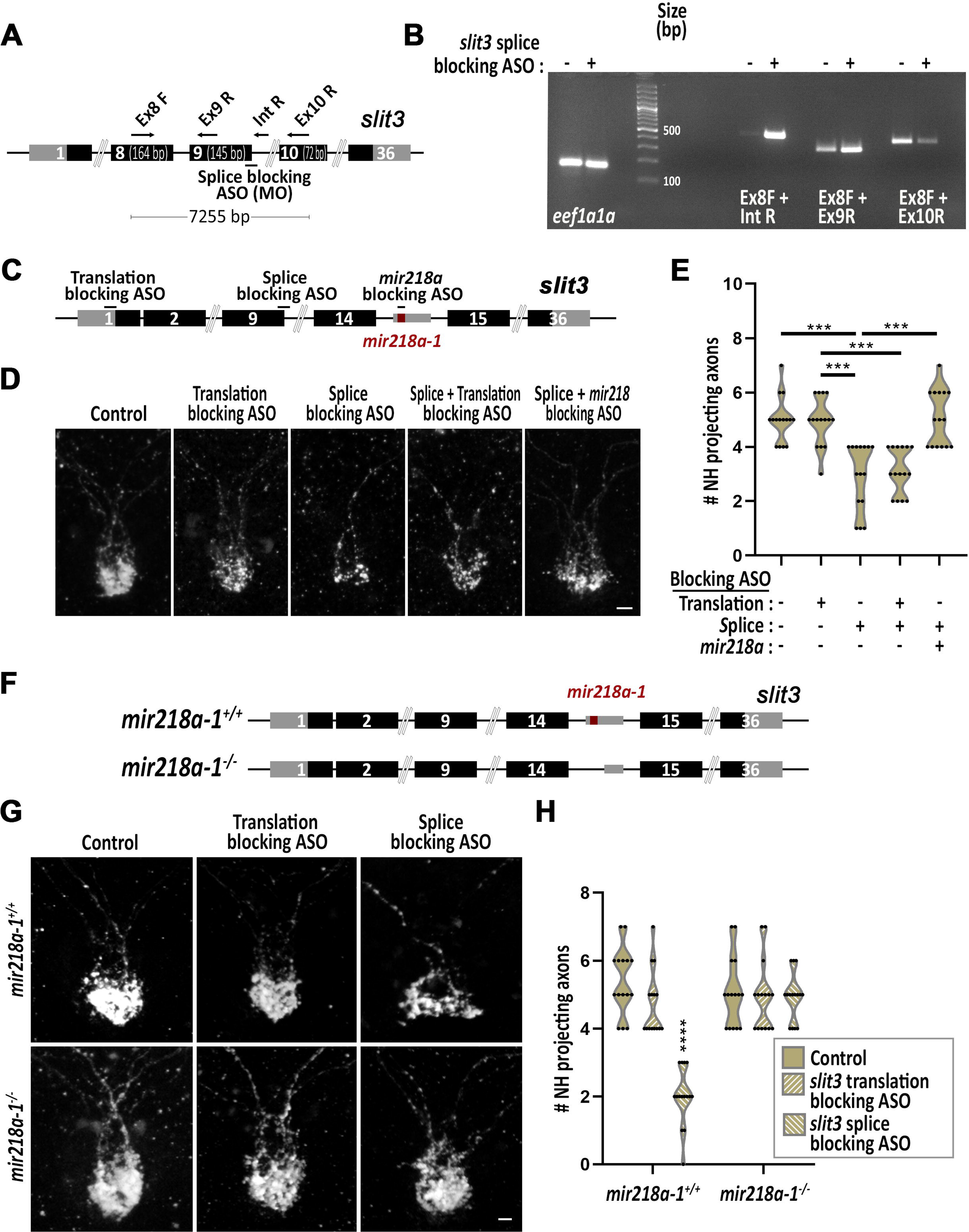
Axonal phenotypes due to *slit3* splice-targeting ASO are due to *mir218a-1*. (**A**) Scheme of *slit3 gene* and approximate location of splice-blocking ASO target and the oligonucleotide primers. (**B**) Gel electrophoresis image showing amplified mRNA splicing products in control embryos and embryos injected with a *slit3* splice-blocking ASOs. Amplicon products of housekeeping gene *eef1a1a* are also shown. The expected splice products and the size of the amplicon products are described in Table S6. (**C**) Scheme of *slit3 gene.* ASO designed to target *slit3* translation, splicing and *mir218a-1* are denoted by black lines and labelled. (**D**) Representative images showing whole-mount confocal microscope imaging of pituitary of 5 dpf transgenic Tg(*oxt*:EGFP) zebrafish following immunostaining with anti-EGFP. Images are from control embryos and following microinjection of individual or co-injected ASOs as listed. Scale 10 µm. (E) Violin plot showing the number of pituitary-projecting OXT axons in 5-dpf control embryos or embryos injected with individual or co-injected ASOs as listed (***p<0.001; Kruskal-Wallis test, n=15). (**F**) Scheme of *slit3 gene* in *mir218a-1* wildtype and knockout zebrafish. (**G**) Representative images showing whole-mount confocal microscope imaging of pituitary of 5 dpf transgenic Tg(*oxt*:EGFP) zebrafish following immunostaining with anti-EGFP. Images are from control larvae and following microinjection of *slit3* translation- or splice-blocking ASOs in *mir218a-1*^+/+^ or *mir218a-1*^-/-^ embryos. Scale 10 µm. (**H**) Violin plot showing the number of pituitary-projecting OXT axons in 5-dpf *mir218a-1*^+/+^ vs *mir218a-1*^-/-^ control embryos or embryos that were injected with *slit3* translation- or splice-blocking ASOs (****p<0.001; Kruskal-Wallis test, n=15).

To prove that *mir218a-1* dysregulation is the cause for the axonal morphogenesis phenotype, we also blocked *slit3* splicing in embryos of previously characterized *mir218a-1*^wz15^ knockout fish (**Fig. 2F**) (Swaminathan et al. 2023). Specifically, we injected *slit3* translation- or splice-blocking ASO in *mir218a-1*^+/+^ and *mir218a-1*^-/-^ single-cell stage embryos. We observed a decrease in pituitary axons in the *mir218a-1*^+/+^ larvae and not in *mir218a-1*^-/-^ larvae (**Fig. 2G and 2H**). Injection of the translation-blocking ASO did not have any effect on the pituitary axonal morphogenesis in neither of the genotype. Our results suggest that, blocking of *slit3* splicing dysregulate *mir218a-1* function and pituitary axonal projections. Previously, studies in zebrafish and mice have shown that *mir218a-1/Mir218-2* embedded in the *slit3/Slit3* introns contain independent *cis*-regulatory elements that decouple the expression of *pri-mir218a-1/pri-Mir218-2* and *slit3/Slit3* transcripts (Punnamoottil et al. 2015; Amin et al. 2021). Overall, our results suggest that blocking of *slit3* splicing potentially leads to override of the cis-regulatory elements-mediated regulation and loss of the decoupling mechanisms that prevents *slit3* and *mir218a-1* co-expression.

### *pank2* and *dnm2a* splice-blocking ASO-induced phenotypes are partially due to intronic *mir103* and *mir199*, respectively

Next, we investigated the generality of the ASO-induced dysregulated miRNA-based phenotypes. Towards this, we chose the *pank2* gene which contain *mir103* in its intron localized between exon 5 and 6. We previously showed that injection of splice-blocking ASOs targeting *pank2* exon1-intron1 and intron1-exon2 junctions in 1-cell stage embryos leads to hydrocephalus and cardiac edema; however, neither a 4-bp nor a 50-bp deletion mutants of *pank2* recapitulated this phenotype (Zizioli et al. 2016; Mignani et al. 2022; Khatri et al. 2020). We hypothesized that the *pank2* splice-blocking-induced phenotype could be partially due to the intronic *miR103*. To test this, we injected 1-cell stage embryos either with control ASO or *pank2* intron1-exon2 junction targeting splice block ASO. In addition, we also co-injected *pank2* splice block ASO with LNA-based *mir103* control ASO or *mir103* blocking ASO (**Fig. 3A**). Unlike the control ASO-injected embryos, majority of the *pank2* splice block-injected embryos developed hydrocephalus and cardiac edema (**Fig. 3B-D** and **G**). Co-injection of *pank2* splice block ASO with *mir103* block ASO led to a partial rescue of the phenotype, whereas co-injection with a control ASO did not result in a partial rescue (**Fig. 3E-G**). The phenotypes were also not due to TP53-based off-target effects as co-injection of *pank2* splice-blocking ASO with *tp53* translation-blocking ASO did not rescue the phenotypes (**Fig. S2A**). These results suggest that the phenotypes resulting from use of *pank2* splice-blocking ASO can be partially attributed to dysregulated intronic *mir103*. In mice, the expression of *Pank2* and *Mir103* genes across various tissues are not correlated, suggesting the presence of cis-regulatory elements that regulate *Mir103* expression or the presence of differential miRNA processing mechanisms (Polster et al. 2010). Our results indicate when *pank2* splicing is blocked, *mir103* expression decoupling mechanisms is potentially overridden.

**Figure 3.**
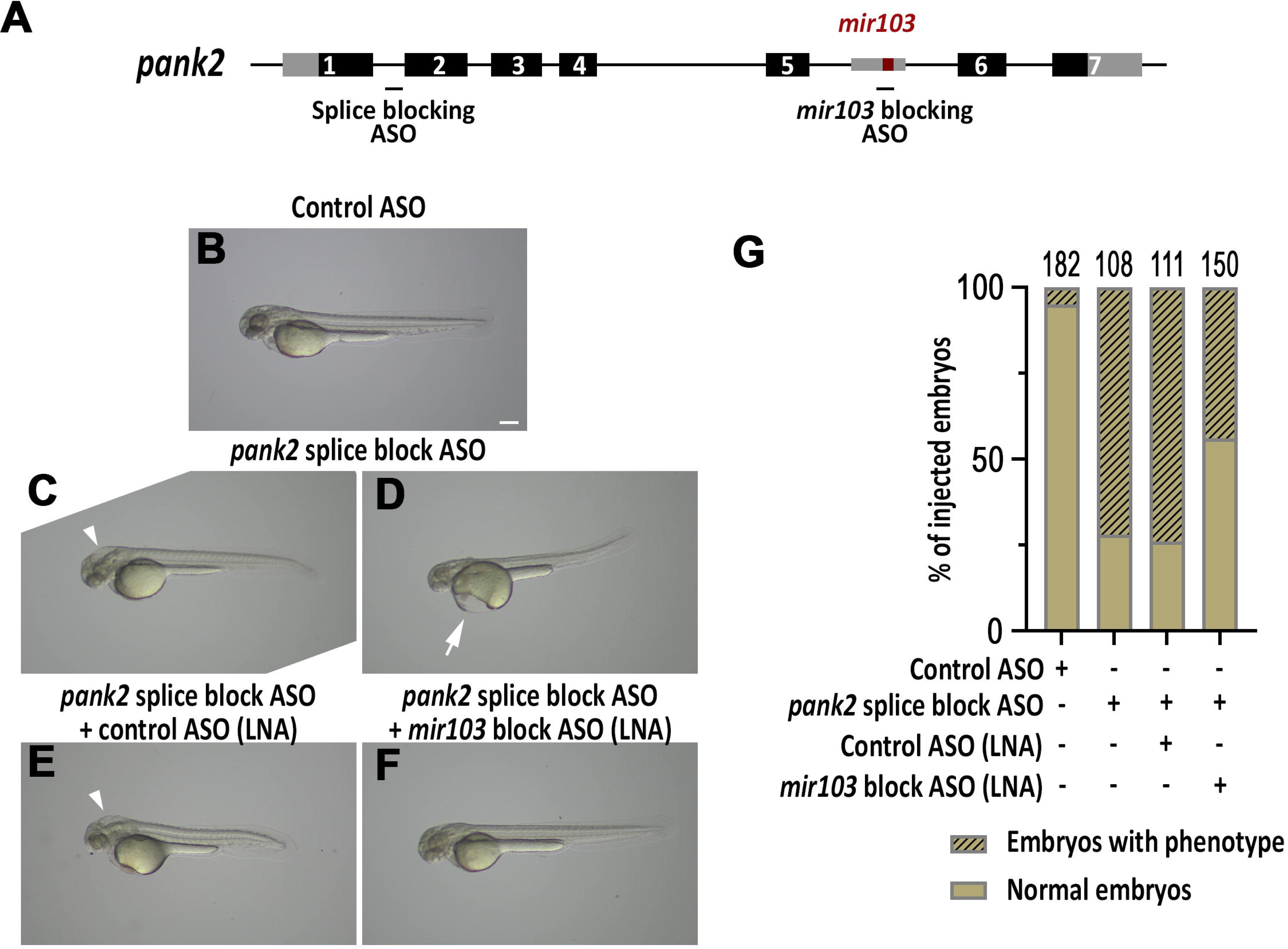
Developmental phenotypes due to *pank2* splice-blocking ASO are partially due to *mir103*. (**A**) Scheme of *pank2* gene. ASOs designed to target *pank2* splicing and *mir103* are denoted by black lines and labelled. (**B-F**) Representative images showing whole-mount light microscope imaging of zebrafish embryos at 2 dpf. Images are from control ASO-injected embryos and following microinjection of *pank2* splice-blocking or *pank2* splice-blocking ASO co-injected with control and *mir103* blocking ASO. White arrows indicate hydrocephalus and white arrow indicate cardiac edema. Scale 250 µm. (**G**) Stacked bar plot showing the analysis of morphological evaluation of embryos performed at 2-dpf and the number of embryos in which the analysis was performed is listed on the top of each bar. Morphological analysis were performed from 3 independent injections. χ^2^=124.7, p<0.0001,df=3.

It is worth to note here that the lack of phenotypes similar to *pank2* splice blocking ASOs in our 4-bp and 50bp *pank2* mutants embryos could be due to genetic robustness or compensation mechanisms (Sztal and Stainier 2020; Bertolaso et al. 2019; Mignani et al. 2022; Khatri et al. 2020). And injection of 1pmol of *pank2* splice-blocking ASO into 50bp *pank2* mutant embryos do not recapitulate the *pank2* splice-blocking ASO phenotypes (Khatri et al. 2020). Hence, our results could be due to *pank2* splice-blocking ASO dose-dependent phenotype. However, it should be noted in ASO-based preclinical trials using mammalian models, higher ASO concentrations are used in order to rescue the phenotype.

Finally, we also investigated if the previously reported *dmn2a* splice-blocking ASO-induced gross developmental phenotypes are also due to its intronic *mir199-2a* that contains duplex miRNAs *mir199-3p* and *mir199-5p* (Gibbs et al. 2013; Bragato et al. 2016). To test this, we injected 1-cell stage embryos with control ASO or exon2-intron2 targeting splice-blocking ASO. In addition, we also co-injected *dnm2a* splice-blocking ASO with LNA-based control ASO or *mir199-5p*-blocking ASO (**Fig. 4A**). We chose to block only *mir199-5p* because it has been shown to be more abundantly expressed than *mir199-3p* (Desvignes et al. 2022). Unlike the control ASO injected embryos, majority of the *dnm2a* splice-blocking injected embryos developed shorter body axis, cardiac edema, and an upward tail curvature (**Fig. 4B-D** and **G**). Co-injection of *dnm2a* splice-blocking ASO with *mir199-5p* blocking ASO led to a partial rescue of the phenotype, whereas co-injection with a control ASO did not result in a rescue (**Fig. 4E-G**). The phenotypes were also not due to TP53-based off target effects as co-injection of *dnm2* splice-blocking ASO with *tp53* translation-blocking ASO did not rescue the phenotypes (**Fig. S2B**). These results suggest that the phenotypes resulting from *dnm2a* splice-blocking ASO can also be partially attributed to dysregulated intronic *mir199-5p*. Studies in mice have shown that the expression of *DNM2* and its intronic *Mir199A1* are positively correlated (Chen et al. 2020). Our results suggest that blocking *dnm2a* splicing dysregulates intronic *mir199-2a* function, but whether this is due to uncoupling of *dnm2a* and *mir199-2a* transcripts and expression is unclear.

**Figure 4.**
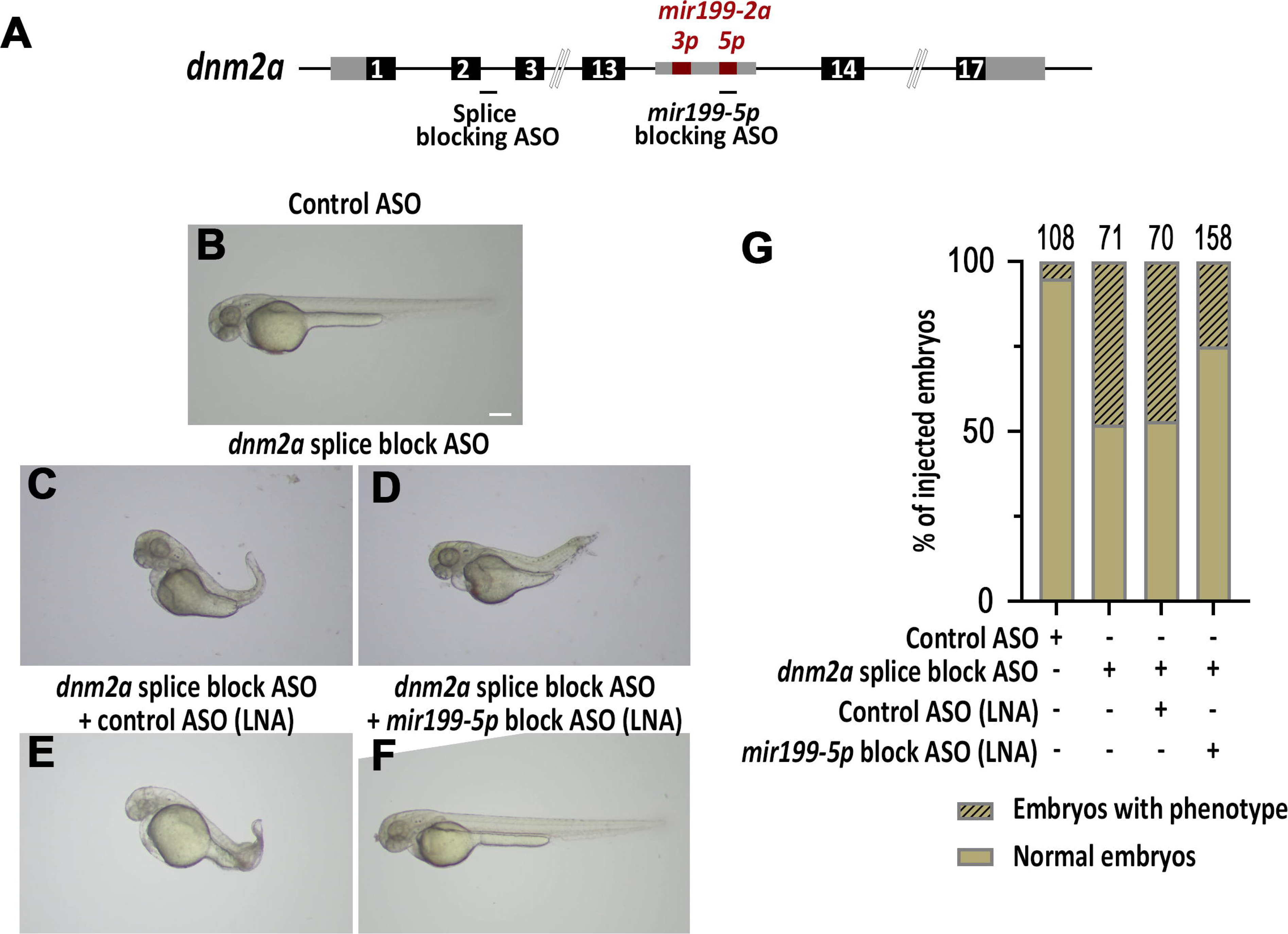
Developmental phenotypes due to *dnm2a* splice-blocking ASO are partially due to *mir199-2a*. (**A**) Scheme of *dnm2a* gene. ASOs designed to target *dnm2a* splicing and *mir199-5p* are denoted by black lines and labelled. (**B-F**) Representative images showing whole-mount light microscope imaging of zebrafish embryos at 2 dpf. Images are from control ASO-injected embryos and following microinjection of *dnm2a* splice-blocking or *dnm2a* splice-blocking ASO co-injected with control and *mir199-5p* blocking ASO. Scale 250 µm. (**G**) Stacked bar plot showing the analysis of morphological evaluation of embryos performed at 2-dpf and the number of embryos in which the analysis was performed is listed on the top of each bar. Morphological analysis were performed from 3 independent injections. χ^2^=58.5, p<0.0001, df=3.

## CONCLUSION

Previous studies using ASO in zebrafish have largely attributed ASO-induced phenotypes that are not reproduced by knockouts to Tp53-based off-target effects (15-20% of all ASO experiments) (Robu et al. 2007; Kok et al. 2015; Eisen and Smith 2008). Another possibility is that the phenotypes could be due to the translated products of the truncated transcripts. However, our data suggest that that there is an additional possible explanation for the observed phenotypes. When *slit3* splicing is blocked, our results indicate that a dysregulated *mir218a-1* causes the phenotype. *mir218a-1* can potentially target and regulate the expression of not only *robo* genes but also any other of the conserved target genes (Amin et al., 2021). We observed similar results also when *pank2* and *dnm2a* splicing was blocked. However, due to the lack of validated anti-Pank2 and anti-Dnm2 antibodies for zebrafish Pank2 and Dnm2a proteins, we were unable to validate the translation-blocking *pank2* and *dnm2a* ASOs. Therefore, we could not exclude the role of potential truncated Pank2 and Dnm2 proteins. Nevertheless, our data contradict the generally held view that off-targets are due to Tp53 and show that phenotypes in 3 different genes tested are at least partially due to the dysregulation of the intronic miRNA encoded by these genes. We are able to rescue the phenotypes (either completely or partially) by blocking the intronic mature miRNA of the targeted gene. Taken together, our work shows that the phenotypes caused by targeted blockade of gene splicing *in vivo* can be attributed to the intronic miRNA (**Fig. 5**).

**Figure 5.**
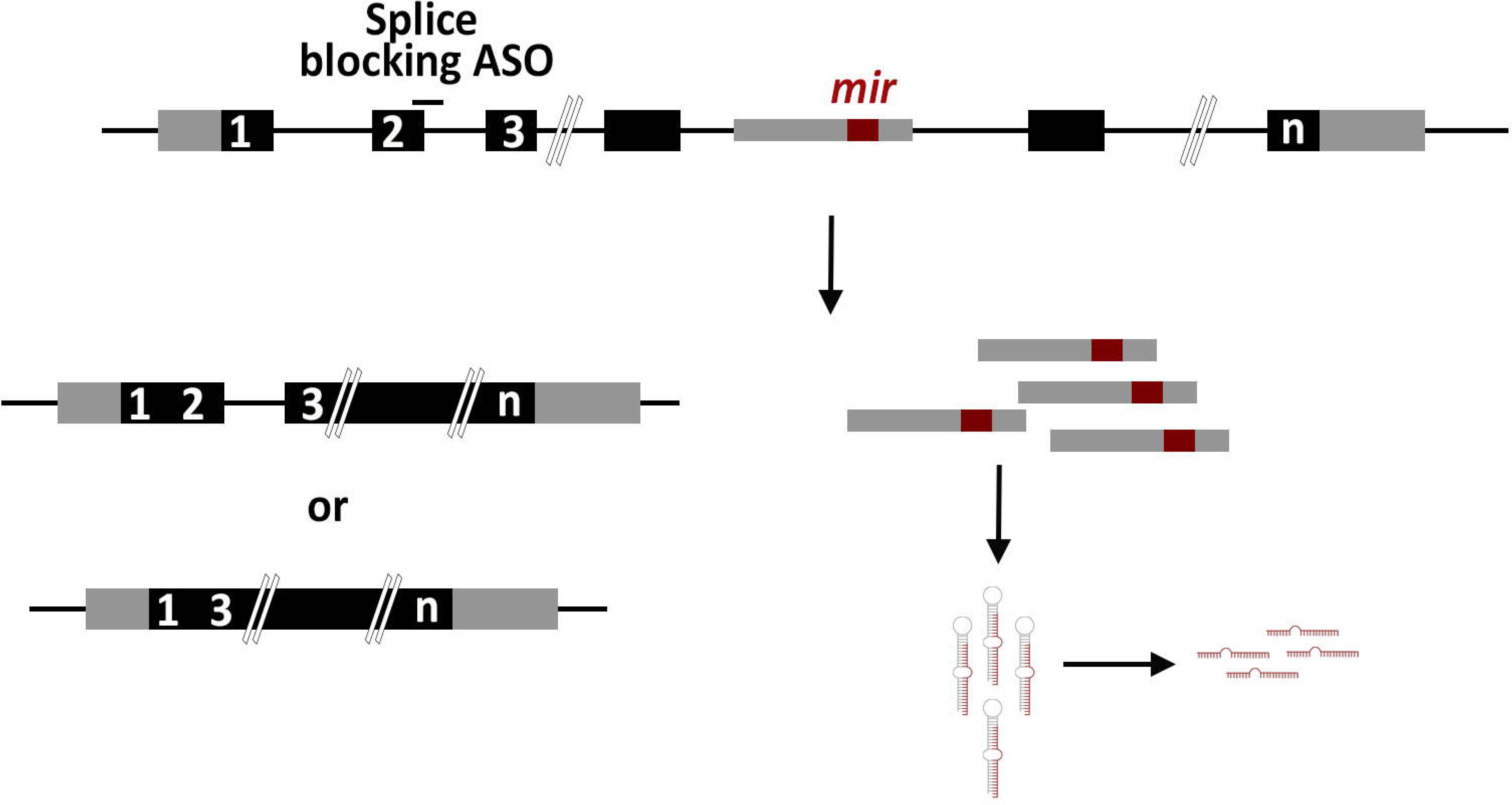
Model. ASO-based targeting of splicing of genes embedded with intronic miRNA may not only lead to intron retention or exon skipping, but also to dysregulation of expression and function of intronic miRNA.

Furthermore, in the case of ASO-induced off-target effects of Tp53 reported in previous studies, the precise reasons for the upregulation of the truncated form of Δ113p53 (Δ113 isoform of p53) induced by ASOs targeting different genes is still unclear (Eisen and Smith 2008). Δ113p53 has been shown to be involved in the DNA damage response by promoting DSB repair pathways (Gong et al. 2015). Pharmacological inhibition of splicing or genetic loss of spliceosomal components has been shown to increase R-loops (RNA-DNA hybrids) accumulation, Tp53 levels, and DNA damage response *in vitro* and in zebrafish (Aguilera and García-Muse 2012; Sorrells et al. 2018; Wan et al. 2015). Whether R-loop formation is a general phenomenon that triggers the expression of Δ113p53 when targeting different genes with splice-blocking ASOs is unknown. However, studies in zebrafish using splicing factor mutants with increased R-loops and Tp53 proteins have shown that the full-length Tp53 is only required for the final apoptotic program and not for mediating the DNA damage (Sorrells et al. 2018). Another possibility is that the miRNA microprocessor (Drosha-DGCR8) complex is "overloaded" with the excess of dysregulated transcripts containing pri-miRNAs. In such cases, the pri-miRNA processing of other transcripts may also be disrupted (Kim and Kim 2007). Whether this is the trigger for Δ113p53 expression is unknown, but Tp53 has been shown to interact with the miRNA microprocessor complex to facilitate pri-miRNA processing (Suzuki et al. 2009). Tp53 interaction with Drosha requires the carboxy-terminal half of the central DNA-binding domain, which is also present in the Δ113p53. Therefore, it is unclear whether the role of ASO-induced expression of Δ113p53 is to promote pri-miRNA processing.

Our *in vivo* zebrafish studies warrant further analysis of the impact of intronic non-coding RNAs when using splice-site targeted ASOs either for therapeutic purposes or for developmental biology studies. Finally, since introns embedded in host genes may contain multiple ncRNAs or classes of ncRNAs, the idiosyncratic phenotypes between translation- and splice-blocking ASOs can potentially be used as a marker to screen for the potential collective role of all the intronic ncRNAs embedded in such host genes. Moreover, with the recent push to move away from animal-based drug safety testing, at least for genes with conserved sequences at the ASO target sites, vertebrate model organisms such as zebrafish embryos must be considered to test the *in vivo* effects of therapeutically promising splice-blocking ASO candidates (Anbalagan 2023).

## MATERIALS AND METHODS

### Bioinformatic analyses

To identify protein-coding genes containing intronic miRNAs and other non-coding RNAs (ncRNAs) in *Danio rerio*, the GRCz11 genome annotation (GCF_000002035.6-RS_2024_08) was analyzed. Specific RNA biotypes, including miRNAs, snoRNAs, and snRNAs, were extracted based on their annotations. Using BEDTools v2.31.1, a sequence of strand-specific intersections was performed to identify ncRNAs overlapping protein-coding genes while excluding those overlapping exonic regions, ensuring only intronic ncRNAs were retained (Quinlan and Hall 2010). GO term analysis on host genes were performed using REVIGO web server (Supek et al. 2011). Adult zebrafish posterior pituitary glial cells expressed genes (mean AMCA+ count >10) were downloaded from (Anbalagan et al. 2018).

### Zebrafish husbandry and maintenance

Zebrafish transgenic lines Tg(*oxt*:EGFP)^wz01^, hybrid wildtype line AB/TL and *mir218a-1*^wz15^ were raised, bred and genotyped according to standard protocols in line with standard procedures and ethical guidelines (Blechman et al. 2011; Swaminathan et al. 2023; Aleström et al. 2020). All experimental procedures were approved by the Local Ethical Committee for Animal Experiments in Poznań. Embryos were grown at 28°C in 0.3X Danieau’s medium at constant uncrowded densities (17.4 mM NaCl, 0.21 mM KCl, 0.12 mM MgSO_4_, 0.18 mM Ca(NO_3_)_2_, 1.5 mM HEPES, pH 7.4).

### Antisense oligonucleotides and primers

The sequences of ASOs and primers can be found in **Table S6**. For ASO-based targeted blocking of gene translation or splicing and mature miRNA function, morpholino and LNA (miRCURY LNA Power Inhibitor) were purchased from Genetools (Philomath, OR, USA) and QIAGEN, respectively. The *slit3, pank2, dnm2,* and *tp53* morpholino-based ASO stock was prepared by dissolving it in distilled water at 1 mM, and one-cell stage embryos were microinjected at 1.67 ng of ASO/embryo. The anti-*mir218*, anti-*mir103* and anti-*mir199-5p* LNA-based ASO stock was prepared by dissolving it in distilled water at 50 µm, and one-cell stage embryos were microinjected at approximately 0.08 ng of ASO/embryo. MO-ASO and LNA-ASO were co-injected at 1.67 ng and 0.08 ng per embryo, respectively.

### In situ hybridization and immunostaining

RNA *in situ* hybridization was performed as described in (Machluf and Levkowitz 2011). For probe preparation, *slit3* (NM_131736.3), *pri-mir218a-1* (host gene NCBI gene ID 80354) and *pri-mir206-1* probes of sizes 346bp, 525bp, and 432bp, respectively were synthesized from a PCR-based templates amplified using the primers listed in **Table S6**.

Immunofluorescence staining was performed as described in (Anbalagan et al. 2019) and using the rabbit anti-EGFP primary antibody (Life technologies/Thermo Fisher, Waltham, MA USA). Secondary antibodies were purchased from Jackson ImmunoResearch Laboratories (West Grove, PA).

### Image acquisition and analysis

Colorimetric *in situ* images were obtained using 5X objective on a Leica DM4B microscope (Leica, Germany). Images of whole embryos fluorescently labeled OXT neurons and axons were obtained by using Nikon A1R confocal microscope with Plan Apo VC 20X objective. Confocal images were analyzed using the FIJI/ImageJ application. The number of oxytocin neurons were counted using the Cell Counter plugin.

### Statistical analyses

Normality tests was performed using Shapiro-Wilks test. ANOVA or Kruskal-Wallis test were used for comparing multiple groups. All data sets were corrected for multiple comparisons. Tukey’s or Dunn’s multiple comparisons tests were used as post-hocs.

## SUPPLEMENTAL MATERIAL

Supplemental material is available for this article.

## ACKNOWLEDGMENTS

We thank Emilia Wysocka, Iwona Kanonik-Jędrzejak and Arleta Kucz for technical and administrative support. We also thank the Zebrafish Core Facility at International Institute of Molecular and Cell Biology in Warsaw for the initial support. S.A. is supported by National Science Centre grants [SONATA-BIS 2020/38/E/NZ3/00090 and SONATA 2021/43/D/NZ3/01798]. H.A. is supported by National Science Centre grant [SONATA 2021/43/D/NZ3/01798].

## AUTHOR CONTRIBUTIONS

H.A. contributed to study design, performed all the experiments, performed data analysis and contributed in manuscript preparation. P.K., W.K., and N.K. performed bioinformatic analysis, assisted in confocal microscopy and in *in situ* hybridization experiments, respectively. N.N. assisted in microinjections. B.G. and D.F. reviewed the manuscript and obtained funding related to *pank2* research. S.A. conceived, designed the study, obtained the funding and wrote the manuscript.

## MAIN FIGURES

**Figure S1. Axonal phenotypes due to *slit3* splice-targeting ASO are not due to OXT neuron loss or *tp53***

(**A**) Graph showing % of zebrafish protein coding genes with and without intronic snRNA and snoRNA.

(**B**) Graph showing the frequency % and no of snRNA per snRNA-embedded protein-coding zebrafish gene.

(**C**) Graph showing the frequency % and no of snoRNA per snoRNA-embedded protein-coding zebrafish gene.

(**D**) Scheme describing the NPO (neurosecretory preoptic area) in 5 days post-fertilization (dpf) transgenic reporter Tg(*oxt*:EGFP).

(**E**) Representative images showing whole-mount confocal microscope imaging of NPO of 5 dpf transgenic Tg(*oxt*:EGFP) zebrafish following immunostaining with anti-EGFP. Images are from control embryos and following microinjection of *slit3* translation- or splice-blocking ASOs. Scale 10 µm.

(**F**) Violin plot showing the number of OXT (oxytocin) neurons in 5-dpf control (n = 14) vs *slit3* translation-blocking ASO (n = 14) and splice-blocking ASO injected embryos (n = 15) (ns indicates not significant; one-way ANOVA).

(**G**) Violin plot showing the number of pituitary-projecting OXT axons in 5-dpf control ASO (n = 19) vs *slit3* splice-blocking ASO injected embryos (n = 17) or embryos coinjected with *slit3* splice-blocking and *tp53* translation-blocking ASOs (n=15) (****p<0.0001; Kruskal-Wallis test).

**Figure S2. Phenotypes due to *pank2 or dnm2* splice-targeting ASO are not due to *tp53***

(**A**) Stacked bar plot showing the analysis of morphological evaluation of embryos performed at 2-dpf in control ASO vs *pank2* splice-blocking ASO injected embryos or embryos coinjected with *pank2* splice-blocking and *tp53* translation-blocking ASOs. χ^2^=91.9, p<0.0001, df=2. The number of embryos in which the analysis was performed is listed on the top of each bar

(**B**) Stacked bar plot showing the analysis of morphological evaluation of embryos performed at 2-dpf in control ASO vs *dnm2* splice-blocking ASO injected embryos or embryos coinjected with *dnm2* splice-blocking and *tp53* translation-blocking ASOs. χ^2^=67.6, p<0.0001, df=2. The number of embryos in which the analysis was performed is listed on the top of each bar

## Supplementary Tables

**Table S1:** List of zebrafish protein-coding genes with intronic miRNAs.

**Table S2:** Gene ontology analysis of zebrafish miRNA embedded host genes.

**Table S3:** List of zebrafish protein-coding genes with intronic snRNAs.

**Table S4:** List of zebrafish protein-coding genes with intronic snoRNAs.

**Table S5:** List of posterior pituitary glial cells expressed intronic miRNA-embedded host genes.

**Table S6:** List and sequence details of ASOs and primers used in the study.

## Supplementary Videos

**Video S1:** Three-dimensional reconstitution of a control 5-dpf old *oxt*:*EGFP* larvae immunostained with anti-EGFP.

**Video S2:** Three-dimensional reconstitution of an anti-EGFP immunostained 5-dpf old *oxt*:EGFP larvae in which *slit3* splice-blocking ASO was injected at 1-cell stage.

