## Supplemental files: Fig S1-S2, Table S1-S6, Video S1-S2. for "Targeted blocking of gene splicing can dysregulate intron-embedded microRNAs": Table S6.docx

**ASO Sequences**

| **Targeted gene** | **Accession number** | **Target** | **Sequence (5’-3’)** | **Reference** | **ASO Type and Source** |
| --- | --- | --- | --- | --- | --- |
| *slit3* | NM_131736.3 | Translation | TATATCCTCTGAGGCTGATAGCAGC | Barresi, M.J. *et al.,* *Development* 2005 | Morpholino from Genetools |
|  |  | Splicing | GAGCTACGGAATCCATACATTTCCA | This study |  |
| *pank2* | NM_001080606.2 | Splicing | CTGTAGTGCAATAATAAGTAGGTGG | Khatri *et al.,* *Bull Exp Biol Med.* 2020 |  |
| *dnm2a* | NM_001030128.1 | Splicing | TGCCGTGCTCATTAACACACTCACC | Gibbs, E.M. *et al.,* *PLoS One* 2013. |  |
| *tp53* | NM_001271820.1 | Translation | GCGCCATTGCTTTGCAAGAATTG | Robu *et al.* 2007, PLoS Genetics |  |
| *Random Control Oligo 25-N* | - | - | NNNNNNNNNNNNNNNNNNNNNNNNN |  |  |
| *mir218a-1* | MIMAT0001868 | Mature miRNA | CATGGTTAGATCAAGCACA | This study | miRCURY LNA Power Inhibitor from Qiagen |
| *mir103* | MIMAT0001816 |  | TCATAGCCCTGTACAA |  |  |
| *mir199-5p* | MIMAT0001277 |  | TAGTCTGAACACTGG |  |  |
| *Negative control A* | - | - | TAACACGTCTATACGCCCA |  |  |

**Primers for probes**

| **Gene name** | **Accession number** | **Fwd (5’ – 3’)** | **Rev (5’ – 3’)** | **Probe size (bp)** |
| --- | --- | --- | --- | --- |
| *slit3* | NM_131736.3 | TAGGATGAGGGAGCGCAAAA | CTCACCACTTCCTGTGAT | 346 |
| *pri-mir218a-1* | host gene  NCBI gene ID 80354 | TCGGGCTTTATATTCCCAGC | AAGTCTGCTTCCCATCAGCC | 525 |
| *pri-mir206-1* |  | AAGAAATACTGCTTTCTGCCCAA | GGTGTGGAGGCTTATGAACTGT | 432 |

**Primers for characterizing *slit3*** (Transcript ID: ENSDART00000146299.3) **splice products**

| **Primer name** | **(5’ – 3’)** |
| --- | --- |
| Ex8 F | ACTCCAACAACCTGCACTGC |
| Ex9 R | GGTTGGCAGGAATCTCAGTCA |
| Int R | ATGACAGGAGGAAGGAGCCA |
| Ex10 R | CTGCTGGGATGTTCTTGATCA |

| **Primer pairs** | **Potential splice products & expected amplicon size (bp)** | |
| --- | --- | --- |
|  | **Control** | **Splice blocking ASO** |
| Ex8 F & Ex9 R | Exon 8 + Exon 9  136 + 124 = 260 | Exon 8 + Exon 9 skipping  none |
| Ex8 F & Int R | none | Exon 8 + Exon 9 + Intron 9 retention  136 + 145 + 118 = 399 |
| Ex8 F & Ex10 R | Exon 8 + Exon 9 + Exon 10  136 + 145 + 38 = 319 | Exon 8 + Exon 9 + Intron 9 retention + Exon 10  136 + 145 + 5895 + 38 = 6214 |
