## Supplementary figures and images for "Targeted blocking of gene splicing can dysregulate intron-embedded microRNAs"

### Fig S1.jpg

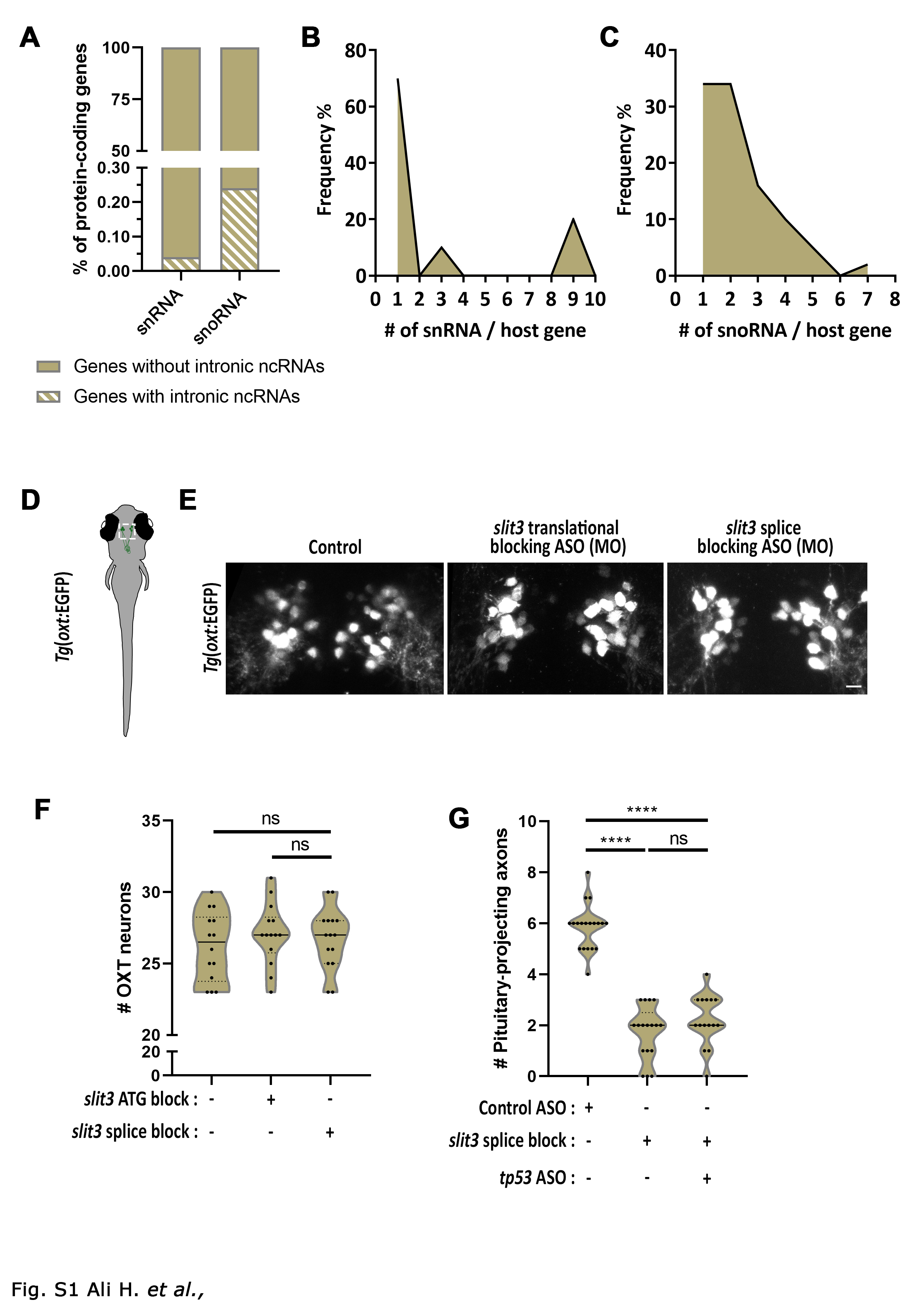

### Fig S2.jpg

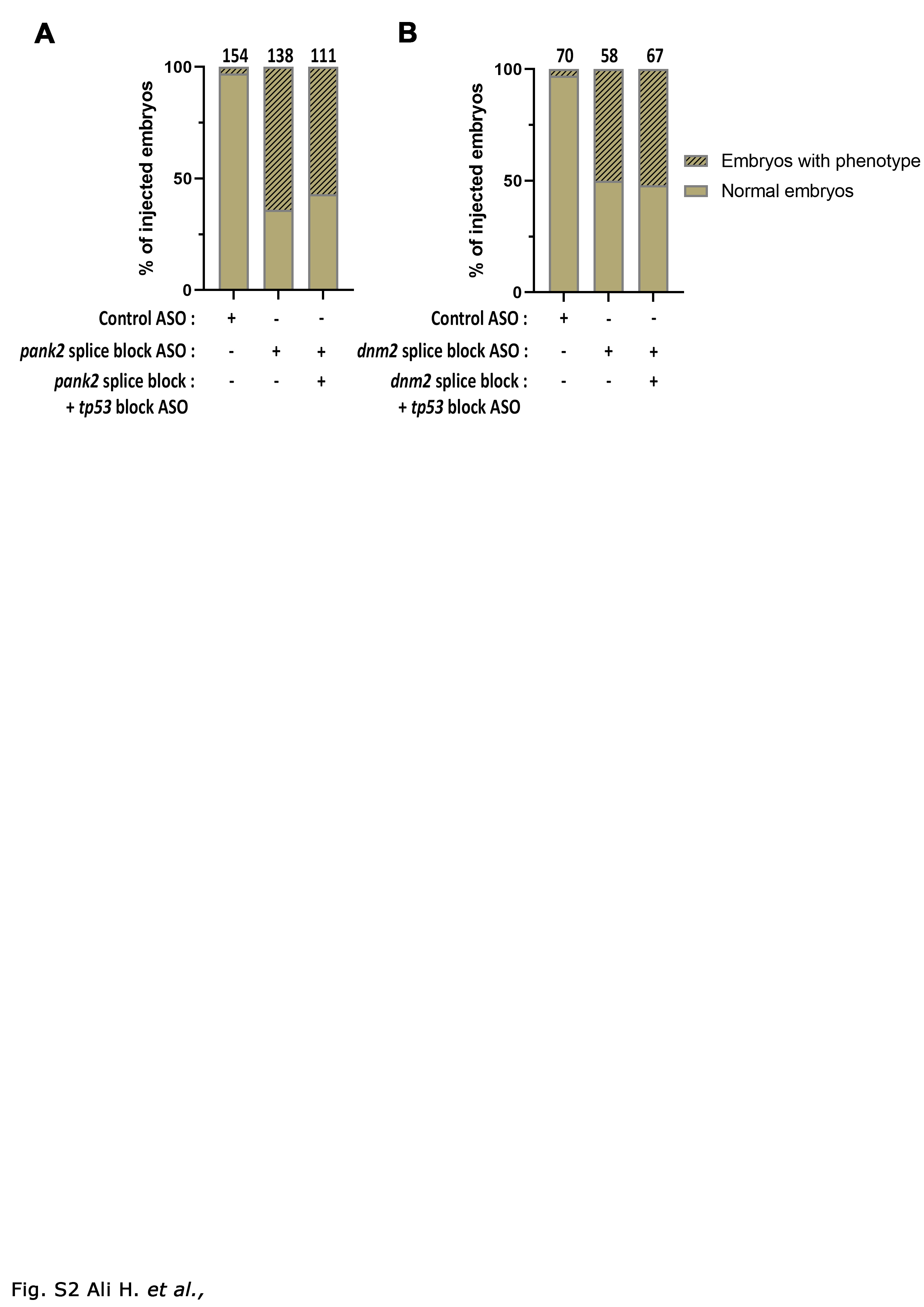
